# RING Finger Protein 11 (RNF11) modulates dopamine release in *Drosophila*

**DOI:** 10.1101/2020.06.29.177857

**Authors:** Eve Privman Champaloux, Nathan Donelson, Poojan Pyakurel, Danielle Wolin, Leah Ostendorf, Madelaine Denno, Ryan Borman, Chris Burke, Jonah C. Short-Miller, Maria R. Yoder, Jeffrey M. Copeland, Subhabrata Sanyal, B. Jill Venton

**Affiliations:** Neuroscience Graduate Program and Medical Scientist Training Program, University of Virginia, Charlottesville, VA; Biogen-Idec, Cambridge, MA; Department of Chemistry, University of Virginia, Charlottesville, VA; Department of Biology, Eastern Mennonite University, Harrisonburg, VA

**Keywords:** RNF11, dopamine, voltammetry, Drosophila, Parkinson Disease, dopamine transporter

## Abstract

Recent work indicates a role for RING finger protein 11 (RNF11) in Parkinson disease (PD) pathology, which involves the loss of dopaminergic neurons. However, the role of *RNF11* in regulating dopamine neurotransmission has not been studied. In this work, we tested the effect of *RNF11* RNAi knockdown or overexpression on stimulated dopamine release in the larval *Drosophila* central nervous system. Dopamine release was stimulated using optogenetics and monitored in real-time using fast-scan cyclic voltammetry at an electrode implanted in an isolated ventral nerve cord. *RNF11* knockdown doubled dopamine release, but there was no decrease in dopamine from *RNF11* overexpression. *RNF11* knockdown did not significantly increase stimulated serotonin or octopamine release, indicating the effect is dopamine specific. Dopamine clearance was also changed, as *RNF11* RNAi flies had a higher V_max_ and *RNF11* overexpressing flies had a lower V_max_ than control flies. *RNF11* RNAi flies had increased mRNA levels of dopamine transporter (DAT) in *RNF11*, confirming changes in DAT. In *RNF11* RNAi flies, release was maintained better for stimulations repeated at short intervals, indicating increases in the recycled releasable pool of dopamine. Nisoxetine, a DAT inhibitor, and flupenthixol, a D2 antagonist, did not affect *RNF11* RNAi or overexpressing flies differently than control. Thus, *RNF11* knockdown causes early changes in dopamine neurotransmission, and this is the first work to demonstrate that *RNF11* affects both dopamine release and uptake. *RNF11* expression decreases in human dopaminergic neurons during PD, and that decrease may be protective by increasing dopamine neurotransmission in the surviving dopaminergic neurons.

## Introduction

Recent work has indicated a role for RING finger protein 11 (RNF11) in Parkinson disease (PD) pathology (Anderson et al., 2007; Pranski et al., 2013b). RNF11 shows homology to E3 ubiquitin ligases and functions as a part of the A20 ubiquitin-editing complex and HECT-type E3 ubiquitin ligase (Kitching et al, 2003). The A20 complex inhibits NF-κB activation in human neurons (Pranski et al., 2012) and persistent NF-κB activation is a hallmark of neurodegenerative disease (Glass et al., 2010; Bellucci et al., 2020). RNF11 variants in the PARK10 locus may increase susceptibility for PD (Lesage and Brice, 2009). RNF11 colocalizes with α-synuclein and Lewy bodies in PD tissue (Anderson et al., 2007). Analysis of genes and extracted brain tissue from Parkinson’s patients revealed *RNF11* gene expression and protein level is decreased in brain tissue affected by neurodegenerative diseases (Li et al., 2014, Pranski et al., 2013b). In rats with the 6-OHDA model of PD, *RNF11* knockdown protected against dopaminergic neuronal cell death and *RNF11* overexpression enhanced toxicity (Pranski et al., 2013a). The enhanced toxicity from *RNF11* overexpression corresponded to downregulation of NF-κB transcribed antioxidants glutathione synthetase and superoxide dismutase 1, anti-apoptotic factor BCL2, neurotrophic factor brain-derived neurotrophic factor (BDNF), and tumor necrosis factor alpha (TNF-α). Therefore the decreased expression of *RNF11* found in surviving neurons is thought to be a compensatory response to PD pathology (Pranski et al., 2013a). This decrease is not correlated with age, making *RNF11* an unchanging candidate as a biomarker of neurodegeneration (Li et al., 2014). While *RNF11* downregulation has been discovered in the context of neurodegeneration, no studies have addressed the effect of RNF11 on neurotransmission.

*Drosophila melanogaster* is a powerful model organism for the study of neurodegeneration and genetic tools exist to study the effects of *RNF11* expression levels on neurotransmission (Hirth, 2010; Shin et al., 2018). A true homolog of *RNF11, CG32850*, has been identified in the *Drosophila melanogaster* genome. This *Drosophila* RNF11 has 87% positive sequence identity with the RING domain of the human protein (Kitching et al., 2003). An A20 ubiquitin-editing complex has not been described in fruit flies. However, homologs for many NF-κB pathway proteins have been found (Khush and Lemaitre, 2000), so it is likely that *Drosophila* RNF11 works in concert with these proteins to edit ubiquitin and inhibit NF-κB function. *Drosophila* is also a good model system for studying dopamine as the dynamics of dopamine release can be measured using fast-scan cyclic voltammetry (FSCV) at carbon-fiber electrodes (Vickrey et al., 2009; Xiao et al., 2014; Privman and Venton, 2015; Pyakurel et al., 2018; Shin and Venton, 2018). FSCV provides information about both release and uptake of dopamine (Vickrey et al., 2013). Thus, dopamine can be measured in genetically altered flies to give a better understanding of how RNF11 affects dopamine release.

The goal of this study was to test the effect of *RNF11* knockdown or overexpression on the release and clearance of dopamine in *Drosophila* larvae. The GAL4/UAS system was used to either overexpress *RNF11* or knock it down using RNAi (Brand and Perrimon, 1993). Dopamine release and uptake were measured with FSCV in the *Drosophila* ventral nerve cord (VNC) (Vickrey et al., 2009). Release was controlled optogenetically using the red light sensitive channelrhodopsin CsChrimson (Privman and Venton, 2015). *RNF11* RNAi flies released double the concentration of dopamine, while *RNF11* overexpressing flies did not release dopamine differently than control. This effect was specific to dopamine, as knocking down *RNF11* in serotonergic or octopaminergic neurons did not alter release. Pulsed stimulations were modeled to obtain the Michaelis-Menten kinetic parameters (Xiao et al., 2014), revealing that V_max_, the maximum rate of dopamine uptake, increased in *RNF11* RNAi flies and decreased in *RNF11* overexpressing flies. These experiments are the first to demonstrate that RNF11 affects the ability of cells to release and uptake dopamine. Thus, the decrease in *RNF11* during PD may lead to increased dopamine neurotransmission, which could be compensating for dopaminergic cell loss.

## Experimental Procedures

### Chemicals

Chemicals and drugs were purchased from Sigma-Aldrich (St. Louis, MO) and solutions were made with Milli-Q water (Millipore, Billerica, MA). Dissections, recordings, and calibrations were performed in a physiological buffer (131.3 mM NaCl, 3.0 mM KCl, 10 mM NaH_2_PO_4_ monohydrate, 1.2 mM MgCl_2_ hexahydrate, 2.0 mM Na_2_SO_4_ anhydrous, 1.2 CaCl_2_ dihydrate, 11.1 mM glucose, 5.3 mM trehalose, pH = 7.4). A 40 mM stock solution of nisoxetine was made in buffer and added to the sample for a final concentration of 20 μM. A 10 mM stock solution of flupenthixol was made in DMSO, diluted in buffer, and added to the sample for a final concentration of 5 μM flupenthixol. For nisoxetine and flupenthixol testing, a pre-drug baseline recording was taken and then the drug was bath applied for 15 minutes before post-drug recordings were taken. Flupenthixol is known to decrease the sensitivity of the electrode for dopamine, so the post-drug data were corrected with a calibration factor to account for the loss in signal (Vickrey and Venton, 2011).

### Electrochemical Measurements

Cylindrical carbon-fiber microelectrodes were made using T-650 carbon fibers (a gift of Cytec Engineering Materials, West Patterson, NJ) as previously described (Jacobs et al., 2011). Electrodes were calibrated *in vitro* using a flow cell to determine the electroactive surface area. The electrodes were exposed to buffer containing 1 μM dopamine, serotonin, or octopamine to determine the response to each neurotransmitter. This calibration factor was used to calculate the amount of neurotransmitter measured *in vivo* as peak oxidation current increases linearly with analyte concentration within the physiological range. Fast-scan cyclic voltammetry was performed using a ChemClamp potentiostat with an n=0.01 headstage (Dagan, Minneapolis, MN), PCI 6711 and 6052 computer interface cards (National Instruments, Austin, TX), and a home built break-out box. TarHeel CV software (a gift of Mark Wightman, University of North Carolina) was used to collect and background subtract the data. Every 100 ms the electrode was scanned from −0.4 to 1.3 V and back at 400 V/s for dopamine detection (Venton and Cao, 2020). For serotonin and octopamine detection, modified waveforms were used to minimize electrode fouling. For the serotonin waveform, the potential was held at 0.2 V, and then scanned up to 1.0 V, down to −0.1 V, and then back to 0.2 V at a rate of 1000 V/s every 100 ms.(Jackson et al., 1995; Borue et al., 2009) For octopamine detection, the potential was held at −0.4 V and scanned to 1.4 V at 100 V/s every 100 ms (Pyakurel et al., 2016). The tissue content of dopamine was measured from 3 pooled larval *Drosophila* ventral nerve cords using capillary electrophoresis coupled with FSCV as described previously (Denno et al., 2015, 2016).

### Fly Strains

Fly stocks were all prepared on a *white*^-^ genetic background. The following fly stocks were used in this study: UAS-*RNF11^JF11021^-*RNAi, UAS-*RNF11^HMJ22085^-RNAi, TH*-GAL4, *Tph*-GAL4, *tdc*-GAL4, *Act5C*-GAL4 (Bloomington stock center), and UAS-CsChrimson (Klapoetke et al., 2014). The UAS-*CG32850* (*dRNF11*) line was generated as follows: an EcoRI/Acc65I fragment of RE41137 (Drosophila Genomics Resources Center, Indiana University), containing a full length cDNA of *Drosophila RNF11*, was ligated into pUASTattB vector and transgenic lines were established by using PhiC31integrase-mediated site specific germ line transformation. Fragments were then cloned into the pENTR Gateway entry clone, followed by LR clonase recombination into the *Drosophila* expression vector pDEST-TWF. Constructs were injected into embryos and progeny screened for positive transformants by Rainbow Transgenic Flies, Inc.

### Tissue Preparation

For genetic expression in dopamine neurons, flies with *TH*-GAL4, UAS-mCD8-GFP were crossed with UAS-CsChrimson (control); UAS-CsChrimson, UAS-*RNF11*-RNAi (*RNF11* knockdown); or UAS-CsChrimson, UAS-RNF11 (RNF11 overexpression). The light senesitive CsChrimson channel was used for optogenetic control of the cells (Klapoetke et al., 2014). For the serotonin experiments, the flies expressed *Tph*-GAL4 instead and for octopamine experiments they expressed *Tdc2*-GAL4. Resulting heterozygous larvae were shielded from light and raised on standard cornmeal food mixed 250:1 with 100 mM all-trans-retinal. The VNC of a third instar wandering larva (either sex) was dissected in PBS buffer. Isolated VNCs were prepared and recorded from as previously described (Borue et al., 2009). The electrode was allowed to equilibrate in the tissue for 10 minutes prior to data collection. A baseline recording was taken for 10 seconds prior to stimulation to facilitate background subtraction.

### Stimulated Neurotransmitter Release

Red-orange light from a 617 nm fiber-coupled high-power LED with a 200 μm core optical cable (ThorLabs, Newton, NJ) was used to stimulate the CsChrimson ion channel. The power output at the fiber end was 0.75 mW and the fiber was centered positioned 75-100 μm from the ventral nerve cord using a micromanipulator. The light was modulated with Transistor-Transistor Logic inputs to a T-cube LED controller (ThorLabs, Newton, NJ), which was connected to the breakout box. TTL input was driven by electrical pulses controlled by the TarHeel CV software, which was used to control the frequency (60 Hz), pulse width (4 ms), and number of pulses (120) in the stimulations.

### Quantitative PCR

For each genotype, 20 adults expressing the Act5C-GAL4 ubiquitous driver were flash frozen and stored at −80 °C before RNA extraction and RT-qPCR analyses. RNA was purified using the RNeasy Mini protocol (Qiagen, Hilden, Germany). Isolated RNA was quantified using a NanoDrop spectrophotometer (Thermo Scientific, Wilmington, DE, USA). 100 microgram of RNA was reverse transcribed using the iScript cDNA Synthesis kit (Bio-Rad, Hercules, CA), according to the manufacturer’s instructions. The cDNA was stored at −80 °C until use.

Primers were designed using primer-BLAST (http://www.ncbi.nlm.nih.gov) (Table 1). RT-qPCR was performed on a CFX Connect Detection System (Bio-Rad, Hercules, CA) using Sso Advanced Universal SYBR Green Mix (Bio-Rad, Hercules, CA). Reactions were carried out in 20 μl total volume, with 200 ng cDNA and 100 nM of each primer. The following amplification program was used in all RT-qPCRs: 95 °C for 2 min, and 40 cycles of 15 s at 95 °C and 15 s at 60 °C. The specificity of each amplified reaction was verified by a dissociation curve analysis after 40 cycles by 2 % agarose gel electrophoresis. Each sample was analyzed in quadruplicate wells and NTC “no-template controls” (without cDNA in the PCR) were included.

The minimum information for publication of quantitative real-time PCR experiments (MIQE) guidelines (Bustin et al. 2009) were used to promote implementation of experimental consistency and to increase the integrity of our results. Serial 10-fold dilutions of each primer pair were used to generate a standard curve and amplification efficiencies (E) were determined based on the slope (M) of the log-linear portion of each standard curve (E = 10-1/M −1). Serial 2-fold dilutions were used to generate a standard curve, and amplification efficiencies (E) were determined based on the slope (M) of the log-linear portion of each standard curve (E = 10^−1/M^ −1 * 100).

### Statistics and Data Analysis

Data are presented as mean +/- standard error of the mean (SEM) displayed as error bars. GraphPad Prism 6 (GraphPad Software Inc, La Jolla, CA) was used to perform all statistics, including one-way and two-way ANOVA with Bonferonni post-hoc tests. Stimulated changes in the extracellular concentration of dopamine were modeled as a balance between discontinuous release and continuous uptake (Wu et al., 2001). A nonlinear regression with a simplex minimization algorithm was used to fit the curves (Wu et al., 2001). The modeling software was stopped when the iteration number continued increasing without a change in the three floated parameters: K_m_, V_max_, and dopamine release per light pulse ([DA]_p_). The goodness of the fit was described using the square regression coefficient (R^2^). The average R^2^ for this work was 0.96 with a standard deviation of 0.02.

## Results

To study the effects of RNF11 on dopamine release, *RNF11* RNAi or overexpression was driven by the tyrosine hydroxylase (*TH*) GAL4 driver, which is a dopamine synthesis enzyme. Dopamine release was stimulated using red light to activate CsChrimson, and extracellular dopamine concentrations were monitored in real-time using FSCV (Privman and Venton, 2015). The pulsed optogenetic stimulation mimics the phasic neurotransmission that is important for reward and motivation in mammals (Venton et al., 2003). Figure 1A shows that knocking down *RNF11* (unless otherwise noted all experiments for RNF RNAi are the *JF11021* allele) caused larger dopamine release than control or *RNF11* overexpression flies. Figure 1B shows example cyclic voltammograms, which confirm dopamine was detected with its characteristic oxidation peak at 0.6 V and reduction peak at −0.2 V. For average peak concentrations, extracellular dopamine concentration varies with genotype (Fig. 1C, one-way ANOVA, p=0.0058) and *RNF11* RNAi flies have significantly larger dopamine release than control, about double the normal concentration (Bonferroni post hoc tests, p<0.01). There was no significant difference between the peak release for control and RNF11 overexpressing flies. To confirm that the increased dopamine release was not specific to the particular RNF11 allele tested, the *RNF11^HMJ22085^* strain was also tested. Similar to the *JF11021* allele, the *HMJ22085* RNAi line also released about twice the amount of dopamine upon activation (Fig. 2).

**Figure 1.**
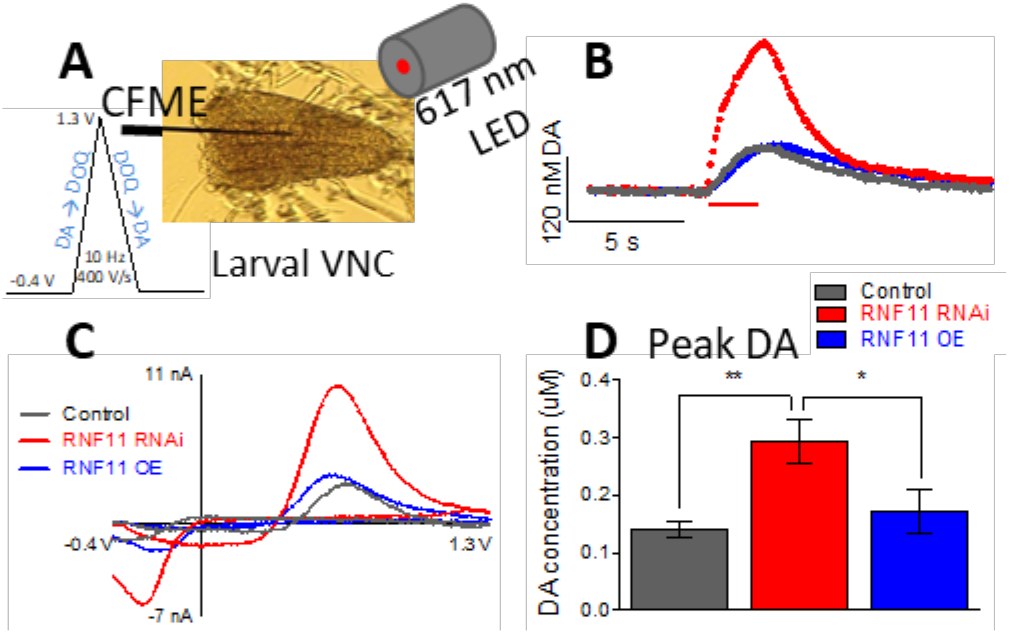
*RNF11* affects dopamine release. (A) Schematic of FSCV recording from larval *Drosophila* VNC. (B) Peak oxidation current of stimulated dopamine in isolated larval ventral nerve cords. The 2 s stimulation is marked by the red line (60 Hz, 120 pulses, 4 ms red light each pulse). The three genotypes plotted are control (grey, *TH*-GAL4; UAS-CsChrimson), *RNF11* knockdown (red, *TH-* GAL4; UAS-*RNF11^JF11021^*-RNAi, UAS-CsChrimson), and *RNF11* overexpression (blue, *TH-* GAL4; UAS-*RNF11*, UAS-CsChrimson). (C) Cyclic voltammograms showing characteristic oxidation peaks for dopamine at 0.6 V and reduction peaks at −0.2 V. (D) Average peak stimulated dopamine concentration changes with genotype (one-way ANOVA, p=0.0019). *RNF11* RNAi flies show greater DA release than control or *RNF11* overexpressing flies (Bonferroni post-test), while *RNF11* overexpression does not affect peak dopamine concentration. Data shown are mean ± SEM (n=10). Asterisks denote significant differences between groups: *p<0.05, **p<0.01.

**Figure 2.**
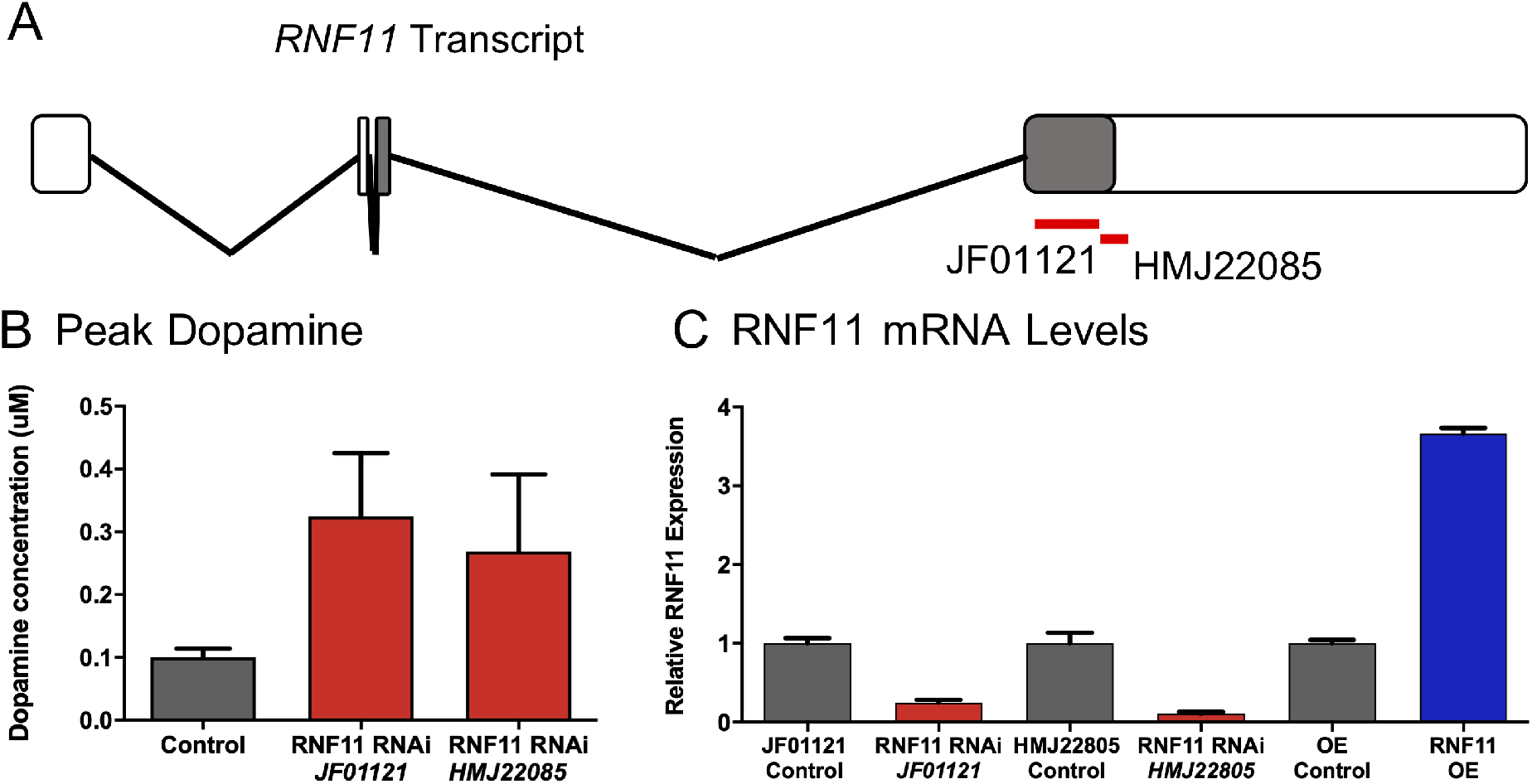
Increased dopamine release is not specific to the RNF11 RNAi allele. (A) The RNAi lines *RNF11^JF11021^* and *RNF11^HMJ22085^* target the 3’ coding region (shaded) of the RNF11 transcript. *RNF11^JF11021^* targets a 300 nt region while *RNF11^HMJ22085^* targets a 21 nt region. The two RNAi alleles overlap only in 3 nt. (B) Average peak dopamine release is significantly higher in both RNF11 alleles (one-way ANOVA, p=0.03), though difference between alleles is insignificant. The three genotypes plotted are control (grey, *TH*-GAL4; UAS-CsChrimson), *RNF11* RNAi^*JF11021*^ (red, *TH*-GAL4; UAS-*RNF11^JF11021^*-RNAi, UAS-CsChrimson), and *RNF11* RNAi^*HMJ22085*^ (red, *TH*-GAL4; UAS-*RNF11^HMJ22085^*-RNAi, UAS-CsChrimson). (C) Capacity of RNF11 knockdown and overexpression were measured by RT-qPCR. The Act5C-GAL4 ubiquitous driver was used to activate RNAi in the *RNF11^JF11021^* and the *RNF11^HMJ22085^* lines. *RNF11^JF11021^* shows a 75% knockdown and the *RNF11^HMJ22085^* shows an 89% knockdown in RNF11 mRNA levels. The UAS-RNF11 overexpression (OE) displays 366% increase in RNF11 levels.

In the present study, *RNF11* was either overexpressed or knocked down with the use of the GAL4/UAS system available in *Drosophila*. To check the overall efficacy of the UAS-*RNF11*-RNAi and the *RNF11* overexpression constructs, the UAS flies were crossed to the universal actin (*Act5C*) driver, and RNF11 mRNA levels were shown to be reduced to 75 ± 4% of control in the RNAi^*JF11021*^ allele. The overexpression of RNF11 by the actin driver increased mRNA levels to 366 ± 8% of control (Fig. 2B). Given the changes in gene expression observed with *Act5C*-GAL4, it is reasonable that similar changes would likewise be observed with other GAL4 driver lines.

To check the specificity of this effect for dopaminergic neurons, *RNF11* RNAi was expressed along with CsChrimson in serotonergic (*Tph*-GAL4) or octopaminergic (*Tdc2*-GAL4) neurons. Neurotransmitter release was monitored using specialized waveforms for those neurotransmitters during a 2 s pulsed red light stimulation. Knocking down *RNF11* in serotonergic neurons did not change stimulated serotonin levels (Figure 3A, t-test, p=0.8054); similarly, knocking down *RNF11* in octopaminergic neurons did not change peak octopamine concentrations (Figure 3B, t-test, p=0.9345). Thus, *RNF11* RNAi selectively increases dopamine release.

**Figure 3.**
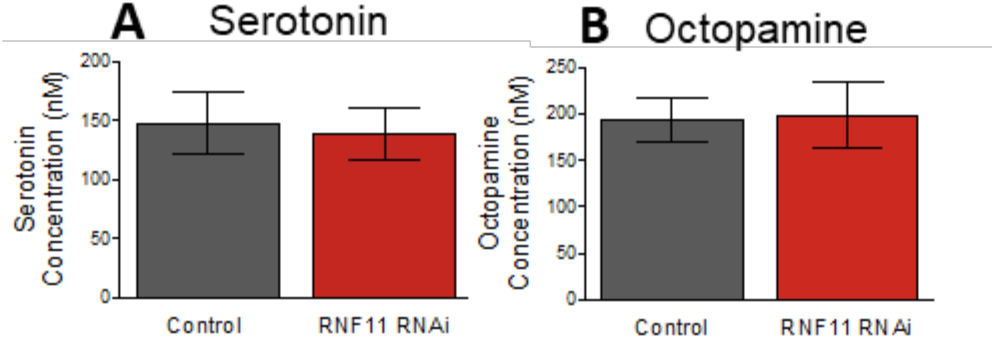
Knocking down *RNF11* with RNAi does not change peak serotonin or octopamine levels. *RNF11^JF11021^* RNAi expressed in (A) serotonergic or (B) octopaminergic cells did not significantly change peak stimulated neurotransmitter levels (unpaired t-tests, p>0.05, n=4-6). Stimulation was 2 s, 60 Hz, 120 pulses. Data are mean ± SEM.

In order to determine the extent to which dopamine release and uptake are affected by *RNF11* knockdown, the current versus time plots (examples shown in Figure 1A) were modeled using Michaelis-Menten kinetics (Figure 4) (Wu et al., 2001). The parameters determined were [DA]_p_, the amount of dopamine released with each pulse of stimulating light, V_max_, the maximum rate of clearance by the dopamine transporter (DAT), and K_m_, the affinity of DAT for dopamine. Dopamine released per pulse of light is affected by genotype (Fig. 4A, one-way ANOVA, p=0.0006) and *RNF11* knockdown significantly increases dopamine release per pulse compared to control (Bonferroni post hoc test, p<0.001). *RNF11* overexpression does not affect [DA]_p_. V_max_ is also significantly affected by genotype (Fig. 4B, one-way ANOVA, p<0.0001). *RNF11* knockdown significantly increases V_max_ compared to control (Bonferroni post-hoc test, p<0.05) and *RNF11* overexpression significantly decreases V_max_ compared to control (Bonferroni post-hoc test, p<0.05). K_m_ is not affected by genotype (Fig. 4C, one-way ANOVA, p=0.3056). Fig. 4D shows the DAT mRNA levels in adult flies where *RNF11* is either knocked down or overexpressed by the ubiquitous *Act5C*-GAL4 driver. There is a main effect of genotype (one-way ANOVA, p=0.0022, n=4) and *RNF11* knockdown produces double the DAT mRNA (p<0.01), while *RNF11* overexpressing flies show similar DAT mRNA levels to controls.

**Figure 4.**
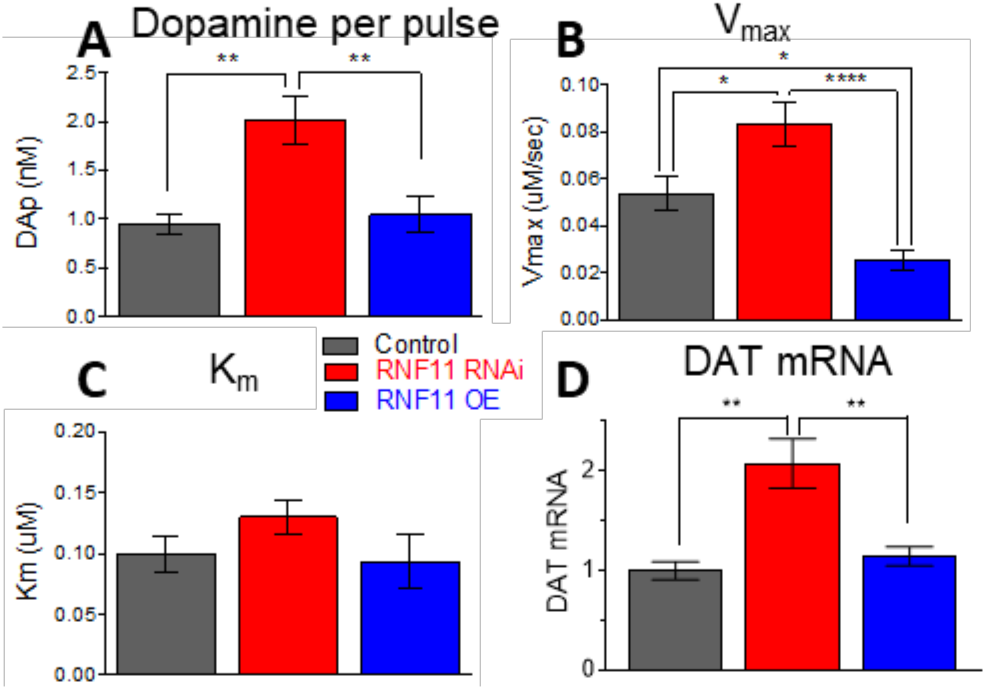
*RNF11* knockdown increases dopamine release and clearance. Dopamine release was modeled and release and uptake parameters determined. (A) Dopamine released per light stimulation pulse ([DA]_p_, 120 pulse, 60 Hz, 4 ms pulse width stimulation) is affected by genotype (one-way ANOVA, n=10, p=0.0006). [DA]_p_ is greater in *RNF11^JF11021^* RNAi flies than control or *RNF11* overexpressing flies (**, Tukey post-test, p<0.01). (B) V_max_ is significantly affected by genotype (one-way ANOVA, n=10, p<0.0001). *RNF11* knockdown significantly increases V_max_ compared to control (*, Tukey post-test p<0.05) and *RNF11* overexpression (****, p<0.0001). *RNF11* overexpression significantly decreases V_max_ compared to control (*, p<0.05). (C) K_m_ is not significantly affected by genotype (one-way ANOVA, n=10, p>0.5). (D) Dopamine transporter (DAT) mRNA levels in qPCR in adults. *RNF11* is driven by the Act5C-GAL4 driver. There is an effect of genotype (one-way ANOVA, p=0.0022, n=4) and *RNF11* RNAi flies have significantly higher DAT mRNA than control flies and *RNF11* overexpression flies (Tukey posttest, **, p<0.01). Data shown are mean ± SEM.

The amount of dopamine release may be due to increases in the total pool of dopamine. Tissue content was measured using capillary electrophoresis coupled to FSCV (Figure 5) (Fang et al., 2011). The dopamine content per larval central nervous system (CNS) was significantly affected by the genotype of the flies (one-way ANOVA, p=0.0003). Post hoc tests showed that the *RNF11* RNAi tissues had significantly more dopamine than the *RNF11* overexpressing tissue (p=0.0002). However, neither the *RNF11* RNAi (p=0.1152) nor the overexpressing (p=0.0510) flies differed significantly from control. This indicates that tissue content did not increase as much as release.

**Figure 5.**
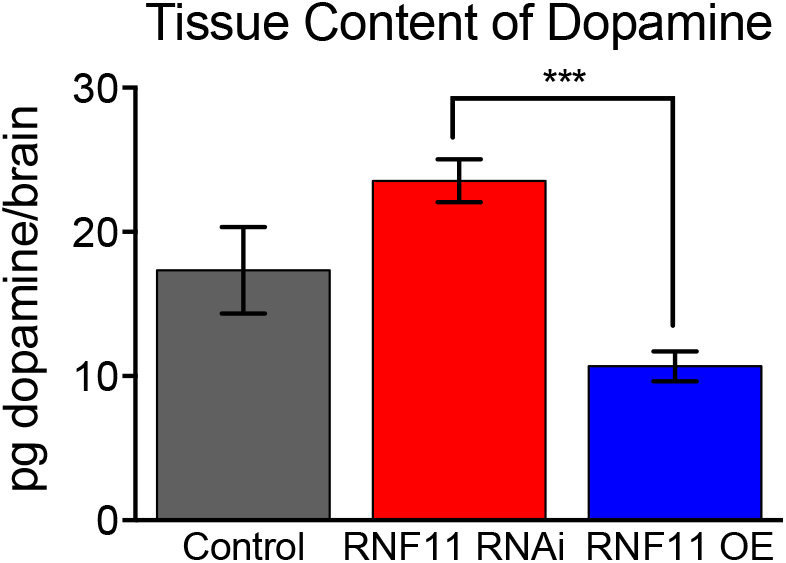
*RNF11* knockdown larval CNS have more dopamine tissue content than *RNF11* overexpressing flies. Capillary electrophoresis coupled to FSCV was used to measure the tissue content of dopamine. The dopamine content was significantly affected by the genotype of the flies (one-way ANOVA, p=0.0003, n=6-9). Post hoc tests showed that the *RNF11* RNAi larval CNS had significantly more dopamine than the *RNF11* overexpressing CNS (p=0.0002). However, *RNF11^JF11021^* RNAi did not differ from control (p=0.1152) nor did *RNF11* overexpressing (p=0.0510). Data shown are mean ± SEM and n=6-9.

To test whether DAT was contributing to release through reverse transport, DAT was competitively inhibited by bath application of 20 μM nisoxetine for 15 minutes (Pörzgen et al., 2001). Figure 5A shows example data of the pre-drug baseline stimulation in an *RNF11* RNAi VNC (orange) and the slowed dopamine uptake after nisoxetine (purple), as evidenced by the increased time to clear dopamine to half the maximum concentration (t_50_). Figure 6B summarizes the data and a two-way ANOVA showed a significant main effect of genotype (p=0.0002) and nisoxetine (p=0.0012), but no interaction. Post-tests show a higher t_50_ for *RNF11* overexpressing flies than both *RNF11* RNAi and control. The percent increase in t_50_ from baseline upon application of nisoxetine was calculated for each genotype, and one-way ANOVA revealed no significant difference (p=0.3558). This indicates that nisoxetine is affecting dopamine uptake equally among the genotypes, despite the slower uptake in *RNF11* overexpressing flies. These results also indicate that the larger dopamine release in *RNF11* RNAi flies cannot be accounted for by reverse transport through DAT.

**Figure 6.**
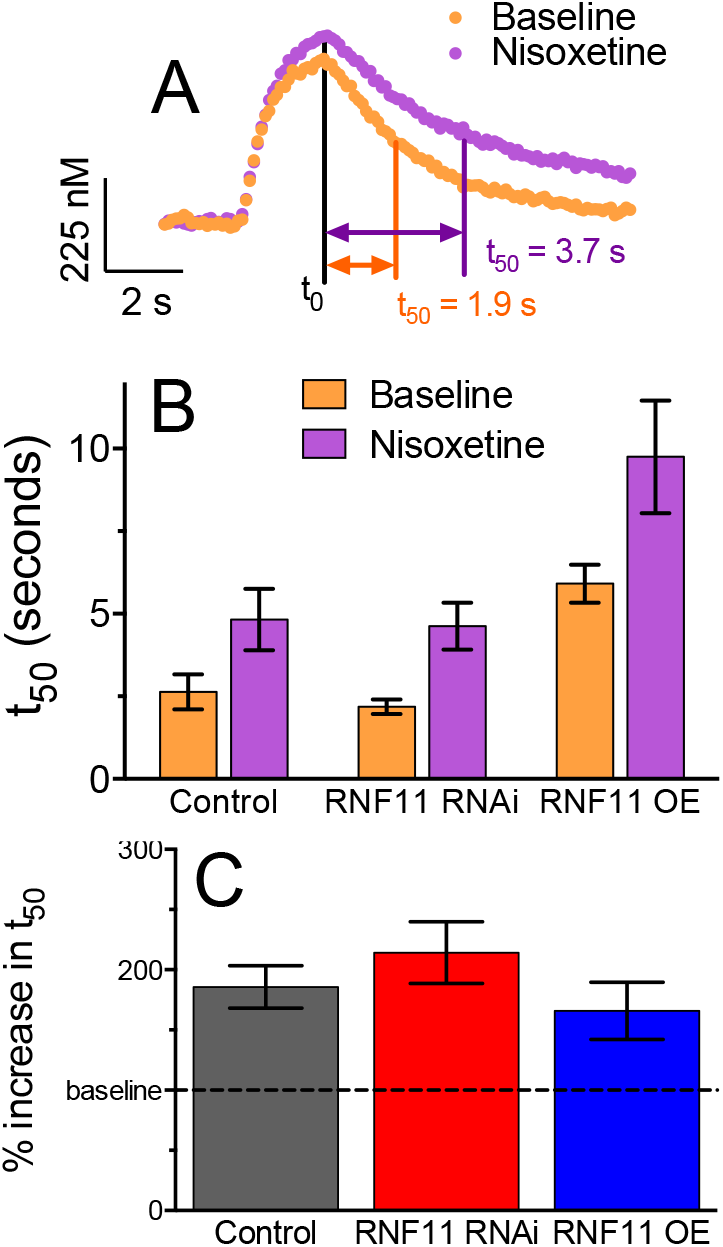
Blocking DAT with nisoxetine causes slowed uptake in all three genotypes. (A) Peak oxidation current for dopamine was monitored over time before (orange) and 15 minutes after (purple) application of 20 μM nisoxetine. This example data shows an *RNF11* RNAi VNC. (B) Nisoxetine caused an increase in t_50_. A two-way ANOVA showed a significant main effect of genotype (p=0.0002, n=4) and nisoxetine (p=0.0012, n=4), but no interaction. (C) The fold increase from baseline upon application of nisoxetine was not significantly different among the different genotypes (one way ANOVA, p=0.3558). Data shown are mean ± SEM and n=4.

Extracellular dopamine levels can be controlled through negative feedback of the D2 dopamine autoreceptor. If *RNF11* knockdown decreases D2 receptor levels in the presynaptic cell membrane, the cell would release more dopamine due to a lack of negative inhibition. D2 was inhibited by bath application of 5 μM flupenthixol for 15 minutes (Vickrey and Venton, 2011). Figure 6A shows example traces of the pre-drug baseline in an *RNF11* RNAi VNC (light green) and the post-drug increased dopamine release (dark green). Figure 6B shows the averaged data. Stimulated dopamine release doubled in all genotypes after flupenthixol bath application. A two-way ANOVA showed a significant main effect of genotype (p=0.0105) and flupenthixol (p=0.0007), but no interaction (p=0.2963). The percent increase in dopamine concentration from baseline upon application of flupenthixol was calculated for each genotype, and one-way ANOVA revealed no significant difference (Fig. 6C, p=0.0659, n=4-7). This indicates that D2 autoreceptors function similarly in all genotypes and *RNF11* knockdown is not increasing dopamine by modulating D2 autoreceptor function.

The turnover of dopamine was assessed by performing multiple stimulations in quick succession. Stimulations were performed every 30 s, which excludes the contribution of dopamine synthesis to the releasable pool since synthesis occurs on the order of 5-10 minutes (Xiao and Venton, 2015). Figure 7 shows that stimulated release decreases with multiple stimulations, but that *RNF11* RNAi flies did not exhibit as large a decrease in evoked dopamine as did control or *RNF11* overexpressing flies. There was a significant main effect of both stimulation number (two-way ANOVA, p<0.0001) and genotype (p<0.0001), but no interaction (p=0.1468). Post hoc tests showed that *RNF11* RNAi flies have a higher plateau than both control and *RNF11* overexpressing flies (p<0.0001 for both). The data were fit to a single-phase exponential decay, which were significantly different (comparison of fits, p<0.0001). Control flies plateaued at 51% of the initial stimulated release, *RNF11* overexpressing plateaued at 59%, and *RNF11* RNAi flies plateaued at 75%. Thus, significantly more dopamine is being recycled for reuse in *RNF11* RNAi flies than in control or *RNF11* overexpressing flies.

**Figure 7.**
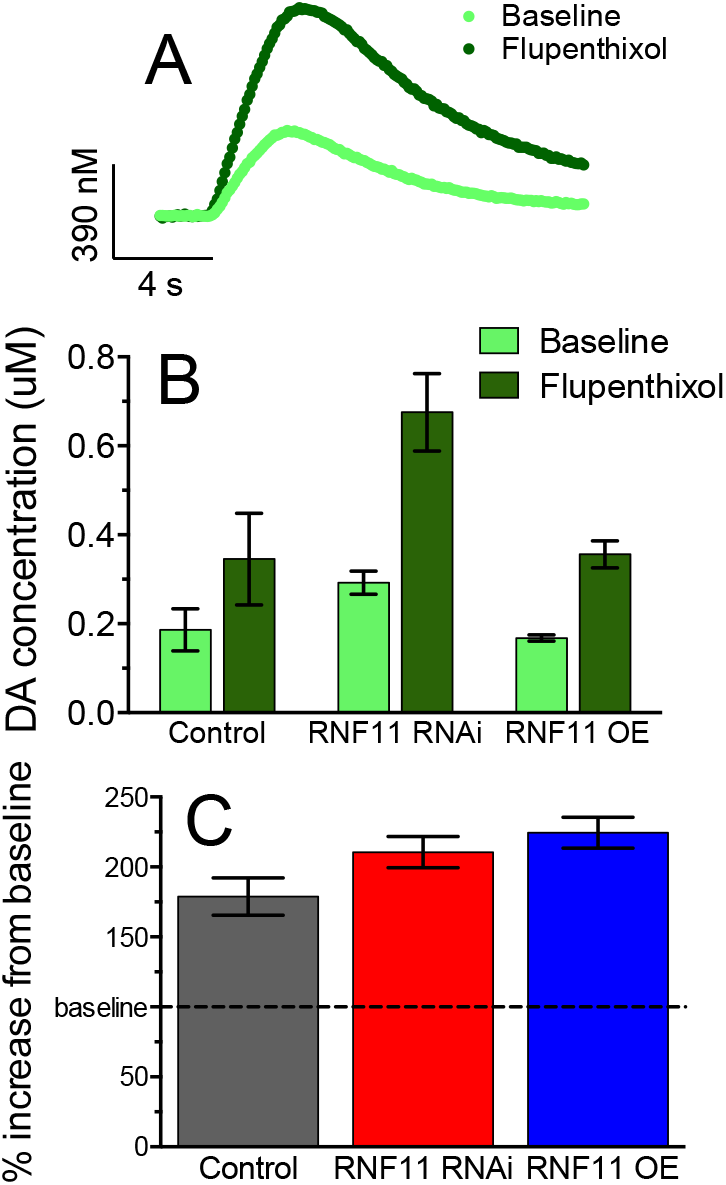
Blocking the D2 autoreceptor with flupenthixol causes increased dopamine release in all three genotypes. Peak oxidation current for dopamine was monitored over time before, 15 minutes, and 30 minutes after the application of 5 μM flupenthixol. (A) Example plot of the effect of 30 minutes of flupenthixol on dopamine release in an *RNF11^JF11021^* RNAi larval CNS. (B) Flupenthixol caused an increase in peak oxidation current of dopamine within 15 minutes in all genotypes. Two way ANOVA revealed a main effect of genotype (p=0.0006) and flupenthixol (p=0.0008), but no interaction (p=0.5800). (C) There was no significant difference among the genotypes in fold increase of dopamine release from baseline upon flupenthixol exposure. (one way ANOVA, p=0.0659, n=4-7).

**Figure 8.**
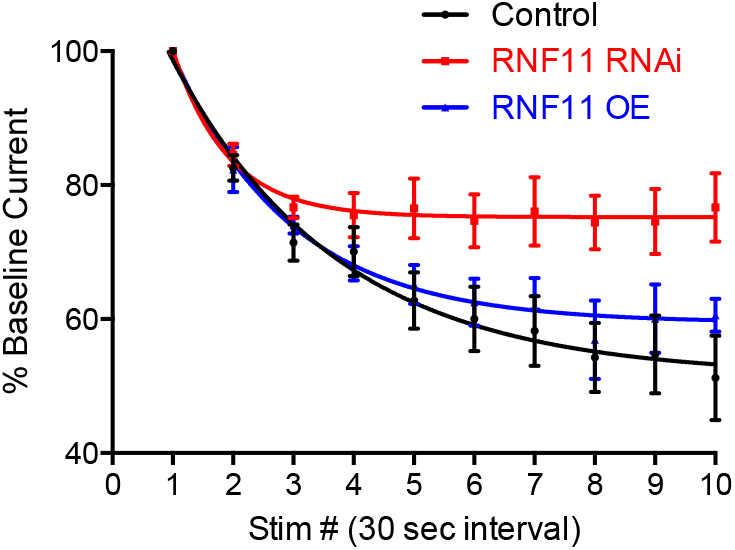
Dopamine turnover is faster in *RNF11* RNAi flies. Larval CNS was stimulated for 2 seconds every 30 seconds, and the peak oxidation current for dopamine was monitored. There was a significant main effect of both stimulation number (p<0.0001) and genotype (p<0.0001), but no interaction (p=0.1468, two-way ANOVA, n=4). Post hoc-tests showed that *RNF11^JF11021^* RNAi flies have a higher plateau than both control and *RNF11* overexpressing flies. The data for each genotype was fit to a single-phase exponential decay, which were significantly different (comparison of fits, p<0.0001).

## Discussion

*RNF11* downregulation has been observed in the late pathogenesis of PD; however early effects of *RNF11* depletion have not been studied. In addition, no study had examined the effect of *RNF11* knockdown on dopamine release and clearance. Here, we describe for the first time that knocking down *RNF11* increases dopamine neurotransmission in *Drosophila* larvae. This effect was specific for dopamine, as knocking down *RNF11* in serotonin or octopamine neurons did not affect stimulated release. *RNF11* RNAi larvae had faster rates of dopamine uptake and more dopamine available for rapidly repeated stimulations than control or *RNF11* overexpressing flies, indicating that *RNF11* knockdown promotes recycling of dopamine and maintenance of the releasable pool. *RNF11* downregulation in neurodegenerative tissue has been postulated as a compensatory response to protect dopaminergic neurons from the insult of PD. Our work indicates that dopaminergic cells with downregulated *RNF11* have more phasic dopamine release. PD patients do not exhibit motor symptoms until a majority of the dopaminergic neurons are dead. Therefore, downregulation of *RNF11* may be a mechanism by which the remaining cells compensate for that loss to maintain the long, symptom-free progression that characterizes this movement disorder.

### *RNF11* knockdown specifically increases dopamine release

*RNF11* RNAi flies had about a quarter of the normal levels of *RNF11* expression and significantly higher stimulated dopamine release than control flies. This doubling in release is about equivalent to blocking the D2 autoreceptors with flupenthixol, which shows that release is greatly affected by *RNF11* knockdown. However, the increase in release is not due to modulation of D2 autoreceptors, since application of flupenthixol doubled the release in *RNF11* RNAi flies as well as controls. One explanation for greater release could be greater synthesis of dopamine. However, *RNF11* RNAi flies did not have significantly higher dopamine tissue content than control flies. Dopamine tissue content in *RNF11* RNAi was significantly higher than *RNF11* overexpressing flies, indicating that synthesis might be somewhat affected by changing *RNF11* levels, but the effect is certainly not a doubling of tissue content. *RNF11* knockdown flies also have greater stimulated release when stimuli were closely applied, showing that they have more dopamine turnover to maintain a larger releasable pool. Thus, the main finding of this work is that *RNF11* RNAi flies are able to increase dopamine release and maintain the releasable pool, and this effect is not primarily mediated by D2 autoreceptors or synthesis.

To test the specificity of release, *RNF11* RNAi knockdown was also performed in serotonergic or octopaminergic neurons. Stimulated release of serotonin or octopamine was not affected by *RNF11* RNAi knockdown. This demonstrates that the effect of *RNF11* knockdown is not universal to all neurotransmitters, but is more specific for dopamine. Dopamine, serotonin, and octopamine are all packaged into vesicles by the same mechanism: vesicular monoamine transporters (VMAT). Humans encode two VMAT genes, *VMAT1* and *VMAT2*, while *Drosophila* has only one, *dVMAT* (Martin and Krantz, 2014). Since *RNF11* RNAi affects only dopamine release, it is unlikely that the mechanism is through dVMAT, since that would also affect other amine neurotransmitters such as serotonin and octopamine.

While *RNF11* knockdown flies had dramatically larger stimulated dopamine release, overexpressing *RNF11* did not significantly change the level of stimulated release. However, the dopamine tissue content of dopamine in *RNF11* overexpressing flies was significantly lower than in *RNF11* knockdown flies and close to significantly lower (p=0.0510) than control flies. These data indicate that there is some compensation happening in the dopamine neurons that allows the *RNF11* overexpressing flies to maintain normal stimulated release, even when the tissue content is somewhat lower. D2 autoreceptors do not appear to be responsible for this effect, as the magnitude of change with flupenthixol was not different than control or *RNF11* RNAi flies. Interestingly, the turnover of dopamine is similar to control, meaning the tissue can support the same level of release with *RNF11* overexpression even though tissue levels are somewhat lower.

In humans, RNF11 functions as part of the A20 ubiquitin editing complex to dampen the signaling cascade that leads to NF-κB activation (Pranski et al., 2013b). Overexpressing *RNF11* leads to a decrease in NF-κB function and knocking down *RNF11* increases NF-κB function (Shembade et al., 2009). In this work we see effects on release, tissue content, and dopamine recycling in dopaminergic neurons with decreased *RNF11*, which may be due to increased NF-κB function. NF-κB has multiple targets and future studies would be needed to discern whether the increase in release works through NF-κB and how its downstream targets work together to affect release. However, we see a limited effect on release and tissue content with *RNF11* overexpression. This may be due to NF-κB already functioning at a low baseline in healthy dopaminergic cells, and further downregulation having limited downstream effects.

### *RNF11* RNAi flies have increased dopamine uptake

*RNF11* RNAi and overexpressing flies also exhibit clear changes in dopamine uptake. *RNF11* RNAi flies had significantly faster uptake than control flies and *RNF11* overexpressing flies had a significantly slower uptake than control. The affinity of the transporter for dopamine (K_m_) did not change. The *DAT* mRNA levels were significantly higher for *RNF11* RNAi flies, but were not significantly lower in *RNF11* overexpression flies. While the qPCR results give a general picture of the amount of *DAT* expressed, the V_max_ corresponds to the number of functioning transporters expressed on the surface. Descrepancies between these two experiments could be due to different rates of internalization, and may explain why measured uptake is slower in *RNF11* overexpression flies while the *DAT* mRNA level is the same. RNF11 may affect the trafficking of DAT and surface expression by modulating the endocytic pathway (Santonico et al., 2010; Gabriel et al., 2013). RNF11 localizes to complexes that mediate endocytosis and endosome trafficking (Pisitkun et al., 2004; Gonzales et al., 2009; Hong et al., 2009; Kostaras et al., 2014), and RNF11 regulates the switch between recycling and degradation of endosomes (Santonico et al., 2014). Therefore, *RNF11* knockdown could lead to a higher proportion of DAT being recycled to the membrane rather than degraded in the endosome pathway. However, there are also NF-κB downstream pathways that could regulate uptake. Overexpressing *RNF11* leads to less activated NF-κB and thus less transcription of brain derived neurotropin factor (BDNF). In mice with 50% of normal BDNF levels, the V_max_ for dopamine uptake was reduced (Bosse et al., 2013), which is consistent with the decrease in V_max_ we observed in *RNF11* overexpressing *Drosophila*. However, *BDNF^+/-^* mice also showed a decrease in dopamine release, which was not found in the *RNF11* overexpression *Drosophila* model. It is important to note that RNF11 has not yet shown to directly assemble ubiquitin chains on any substrate but itself, and in fact might act as an adapter for other E3 ligases (Santonico et al., 2015; Budhidarmo et al., 2018). Thus, the RNF11 effects on release and uptake cannot be explained by just one of the known downstream targets of NF-κB and certainly involves a more complicated protein machinery.

The turnover of dopamine in *RNF11* RNAi flies is also faster, indicating that the mechanisms for dopamine repackaging and release are upregulated when *RNF11* is knocked down. The increased uptake would lead to more dopamine being available for recycling, thus dopamine release is maintained for closely repeated stimulations. Previous studies have shown that the releasable pool of dopamine is dependent on transporters in this time frame,(Xiao and Venton, 2015) and the finding of faster uptake in *RNF11* RNAi flies is consistent with this being the main mechanism of releasable pool regeneration. This maintenance of the pool is not explained by the reverse transport of dopamine through DAT or the downregulation of D2 dopamine autoreceptors. There is also a decrease in the uptake of dopamine in *RNF11* overexpressing flies, allowing dopamine to be active in the extracellular space for longer. Release and turnover levels are maintained in *RNF11* overexpressing flies, despite the lower tissue content and slower uptake. This normal maintenance of the releasable pool points to other mechanisms that are helping to maintain release in *RNF11* overexpressing flies and future studies could probe the possible mechanism of the regulation of dopamine turnover. The increased tissue content, increased release, and faster turnover of dopamine seen in *RNF11* RNAi flies could also occur due to a decreased metabolic breakdown of dopamine in the cytoplasm after uptake. The dopamine metabolic pathways are poorly characterized in *Drosophila*; however they would also likely metabolize serotonin and octopamine (Suh and Jackson, 2008; Freeman et al., 2013; Ueno and Kume, 2014; Yamamoto et al., 2014). Therefore, metabolism is unlikely to be the cause of the dopamine release and turnover effects upon *RNF11* knockdown.

### Implications of RNF11 effects on dopamine neurotransmission in PD

RNF11 affects many pathways important to proper neuronal functioning that could make it important in the cellular response to PD. While dopamine neurons die in early PD, there is a prolonged compensation period, during which remaining neurons continue to support dopaminergic neurotransmission (Hawkes et al., 2010). *RNF11* is downregulated in neurodegenerative diseases, including PD (Pranski et al., 2012, 2013b). Here we present the first evidence that *RNF11* knockdown increases dopamine release and uptake, which may represent a mechanism by which dopaminergic cells maintain functional levels of dopamine despite the loss of a substantial percentage of the local neuronal population. Future studies could examine the importance of RNF11 in PD by simultaneously manipulating *RNF11* and known PD genes in *Drosophila*, such as *PINK1* or *parkin* (Park et al., 2006) to see the phenotypic effects of *RNF11* on disease progression as well at the effects on dopamine neurotransmission. In addition, studies to change the levels of *RNF11* knockdown could examine the extent to which further decreases in *RNF11* increase dopamine release and whether *RNF11* could be a target for future PD therapies.

## Conclusions

*RNF11* has been implicated in the pathophysiology of human PD, but its effect on dopamine neurotransmission was unknown. Here, in *Drosophila*, we show that knocking down *RNF11* doubled dopamine release, while *RNF11* overexpression did not change release. This effect was dopamine specific, as *RNF11* knockdown does not have a similar effect on the serotonergic or octopaminergic neurotransmitter pathways. *RNF11* knockdown increased, and *RNF11* overexpression decreased the rate of dopamine uptake, showing that *RNF11* affects both dopamine release and uptake. Future studies could identify the extent to which the effect of *RNF11* is mediated by NF-κB or endosome recycling. These results are important for understanding PD, as the upregulation of dopamine signaling may help maintain dopamine function in PD patients that have reduced *RNF11* expression. *RNF11* may also be a future biomarker or drug target for detecting or treating PD.

## Abbreviations

BDNF: Brain derived neurotrophic factor
[DA]_p_: Dopamine release per light pulse
DAT: Dopamine transporter
PD: Parkinson disease
RNF11: Ring finger protein 11
TH: Tyrosine hydoxylase
VMAT: Vesicular monoamine transporter
VNC: Ventral nerve cord

## Acknowledgments

The authors would like to thank the Gaultier, Hirsh, and Jayaraman labs for reagents and fly stocks and Karol Cichewicz for his help with fly selection. Work was funded by NIH R01MH085159 and by Biogen-Idec.

